# *Streptococcus pyogenes* capsule promotes microcolony-independent biofilm formation

**DOI:** 10.1101/609065

**Authors:** Artur Matysik, Kimberly A. Kline

## Abstract

Biofilms play an important role in the pathogenesis of Group A Streptococcus (GAS), a gram-positive pathogen responsible for a wide range infections and significant public health impact. Although most GAS serotypes are able to form biofilms, there is large heterogeneity between individual strains in biofilm formation, as measured by standard crystal violet assays. It is generally accepted that biofilm formation includes initial adhesion of bacterial cells to a surface, followed by microcolony formation, biofilm maturation, and extensive production of extracellular matrix that links together proliferating cells and provides a scaffold for the three-dimensional biofilm structure. However, our studies show that for GAS strain JS95, microcolony formation is not an essential step in static biofilm formation, and instead, biofilm can be effectively formed from slow-growing or non-replicating late exponential or early stationary planktonic cells, via sedimentation and fixation of GAS chains into biofilms. In addition, we show that the GAS capsule specifically contributes to the alternative, sedimentation-initiated biofilms. Microcolony-independent, sedimentation biofilms are similar in morphology and 3-D structure to biofilms initiated by actively dividing planktonic bacteria. We conclude that GAS can form biofilms by an alternate, non-canonical mechanism that does not require transition from microcolony formation to biofilm maturation, and which may be obscured by biofilm phenotypes that arise via the classical biofilm maturation processes.

**IMPORTANCE:** The static biofilm assay is a common tool for easy biomass quantification of biofilm forming bacteria. However, *S. pyogenes* biofilm formation as measured by the static assay is strain dependent and yields heterogeneous results for different strains of the same serotype. In this study, we show that two independent mechanisms, for which the protective capsule contributes opposing functions, may contribute to static biofilm formation. We propose that separation of these mechanisms for biofilm formation might uncover previously unappreciated biofilm phenotypes that may otherwise be masked in the classic static assay.

## INTRODUCTION

*Streptococcus pyogenes* (Group A Streptococcus (GAS)) is a human pathogen responsible for a variety of disease states, ranging from superficial infections such as pharyngitis to severe infections such as necrotizing fasciitis, with a global impact on mortality and morbidity [1]. Biofilm formation is thought to be an important GAS virulence factor, and many GAS strains can form biofilm *in vitro* [1-3]. Biofilm-like GAS communities have been observed in tonsillar reticulated crypts, suggesting a role for biofilm in asymptomatic GAS carriage [4]. Increased antibiotic tolerance of GAS biofilms has been also proposed as an important reason for antibiotic treatment failure [2, 5]. However, the ability to form biofilm is often strain dependent, and can be heterogeneous even for isolates belonging to the same serotype. Differential regulation of primary adhesin expression, aggregation tendency, or yet unknown alternative mechanisms responsible for biofilm maturation have been proposed to explain this variability [1, 3].

Static and flow cell biofilm assays are the two most commonly used methods to study biofilm development in GAS and other organisms. Flow cell biofilm assays have the advantage of biofilm growth in the absence of planktonic cells, in a nutrient-rich and metabolite-poor environment, enabling continuous and detailed observation of biofilm maturation over time. However, flow cell biofilms are labor-intensive to assemble and perform, limiting the variety of conditions that can be screened in a quantitative way [6, 7]. By contrast, biofilm growth under static conditions, typically monitored in 24-well polystyrene plates followed by crystal violet or safranin staining, is a simple, high throughput method to quantify overall biofilm biomass. A drawback of the static biofilm assay is nutrient depletion and metabolite accumulation over time, both of which can affect biofilm maturation [8]. Static biofilm assays are strongly influenced by primary cell to surface interactions, rendering analysis of the entire biofilm life cycle more difficult [8, 9]. Despite these limitations, static biofilm assays remain the major tool for quantitative assessment of GAS biofilm formation [3, 8, 10-15]

Several recent studies by us and others have implicated biofilm formation with necrotizing fasciitis (NF) [16-19]. In this study, we sought to dissect the mechanism of biofilm formation of the GAS NF-associated strain JS95. Modifying the commonly used static biofilm assay, we discovered that two parallel mechanisms contribute to GAS biofilm formation: one that proceeds through a classic microcolony proliferation-dependent stage and another that is seeded via proliferation-independent sedimentation. Capsule has been reported as a GAS biofilm factor, but its precise contribution to biofilm development has been unclear [1, 13]. Using the modified static biofilm assay developed here, we demonstrate opposing contributions of the capsule in each biofilm development mechanism.

## RESULTS

### GAS JS95 biofilm *in vitro*

We previously described determinants that are important for necrotizing GAS strain JS95 (M14 serotype) biofilm formation in association with host cells and therefore grew GAS in DMEM cell culture medium in that study [17]. Here we used the more commonly used GAS growth medium of Todd-Hewitt broth with 2% yeast extract (THY) supplemented with 0.5% glucose, which was previously demonstrated to improve adherence and biofilm formation [1, 13, 15, 20]. Biofilms were assayed in 24-well polystyrene plates following static incubation at 37°C for 24 hours. Although some GAS strains require additional surface coating (e.g. ploy-L-lysine, collagen, fibronectin, fibrinogen) for GAS attachment and biofilm formation [3, 13], JS95 formed biofilm on polystyrene plates without a need for additional surface treatment. Confocal microscopy showed extensive three-dimensional biofilm structure **(Figure 1a),** which is typical for most GAS biofilms [3]. As expected, we also observed a similar biofilm architecture by scanning electron microscopy (SEM) **(Figure 1b).** Although biofilm matrix-like structures have been observed by SEM in some GAS biofilm studies [3, 21], we did not observe similar structures. An absence of visible EPS matrix in SEM images of GAS biofilms has been previously reported for serotypes M6, M18, M49 [3], whereas matrix-like material has been observed in MGAS5005 biofilms [21]. The heterogeneity in visible biofilm matrix may be due to differences in sample preparation, strains differences, or other experimental variations. Wheat-germ agglutinin (WGA), a lectin which binds carbohydrate-containing extracellular polymeric substances (EPS), has been used as a marker of biofilm matrix for GAS and other organisms [16, 22, 23]; however, WGA staining did not reveal extracellular biofilm matrix for strain JS95 **(Figure 1c).** We also confirmed that the apparent cell-associated WGA staining is not due to capsule staining by WGA as reported in some studies [16], since the *hasA* capsule mutant showed an identical staining pattern **(Figure S1).** Hence, these findings show that GAS strain JS95 forms 3D structures consistent with biofilm architecture described for other GAS strains, but we were unable to visualize extracellular biofilm matrix.

**Figure 1.**
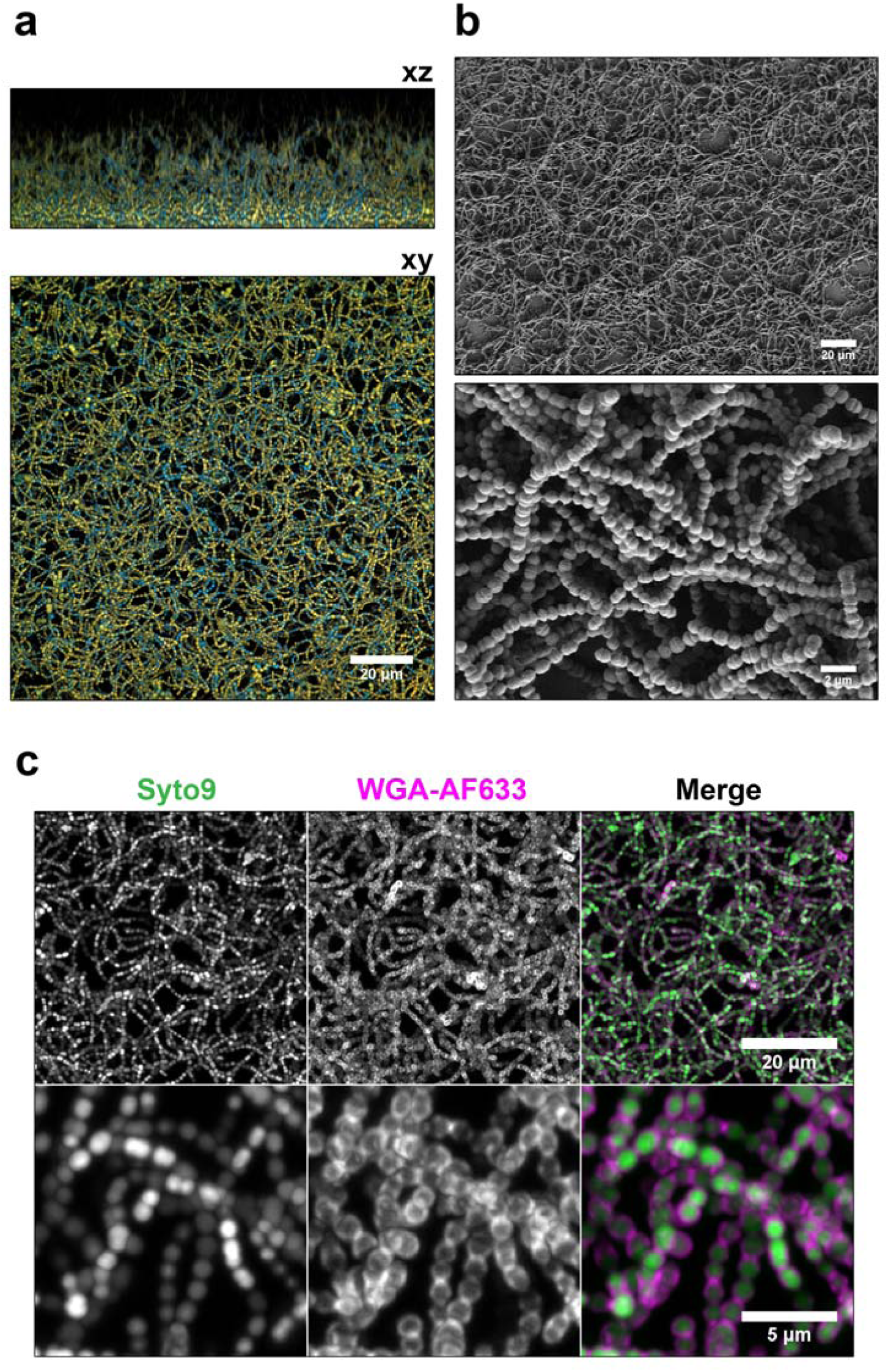
GAS JS95 biofilm morphology. **(a)** JS95 biofilm was grown on polystyrene plate stained with Hoechst33342 (blue), Syto9 (green), and propidium iodide (red). Orange colour represents dead cells. A representative color-merged volume projection of a Z-stack is shown. **(b)** Scanning electron micrographs of JS95 biofilm grown on polystyrene at magnification of 500× (top) and 10,000× (bottom). **(c)** Maximum projection of JS95 biofilm Z-stacks stained with Syto9 (dsDNA) and WGA-Alexa Fluor 633 (carbohydrate/EPS).

### GAS can form biofilms from early stationary planktonic culture

Biofilm development is generally thought to initiate by cell adhesion to a surface followed by the formation of microcolonies, extensive extracellular matrix production, and proliferation, together resulting in a mature biofilm structure [1, 9, 24]. Typically, GAS static biofilm assays are conducted by dilution of overnight cultures into fresh growth medium and incubation in 24-well polystyrene plates without agitation for several hours prior to washing and staining for biofilm biomass [8, 13, 16, 20, 25-27]. Several GAS adhesins, such as M-protein, collagen-like surface protein, and fibronectin-binding protein are upregulated in early stages of planktonic growth [28], and are thought to be important for static biofilm formation [3]. We therefore examined biofilm formation at different stages of planktonic growth, predicting that biofilms initiated from later stages of planktonic growth would exhibit attenuated biofilm formation, due to reduced adhesin expression [28] and reduced proliferation. To test this hypothesis, we designed a simple planktonic transfer assay, in which the same volume of planktonic GAS culture grown in a conical tube (with occasional agitation to prevent biofilm formation and sedimentation) was transferred into a 24-well plate (1ml/well) at various times throughout the growth curve. Therefore, although fewer cells were transferred at early timepoints, the medium contained nutrients allowing for continued growth and biofilm formation. By contrast, at later timepoints, despite a high number of cells being transferred, the medium lacked nutrients supporting further growth. We measured the optical density (OD600) and determined the CFU at each time point to monitor growth rate and planktonic cell viability prior to transfer. 24-well plates were then incubated for 24 hours and the resulting biofilm biomass was quantified using crystal violet (CV) **(Figure 2a)**. Based on this analysis, biofilms were initiated from planktonic cultures at time points approximately representing the following stages of growth: inoculation (0hrs), early exponential (2hrs), mid exponential (4hrs), late exponential (6hrs), early stationary (8 hrs), and later stationary (10 and 12 hrs). CFU enumeration performed at the time of transfer peaked at 6hrs (2.4 x 10^8^ CFU/ml), then decreased to 4 x 10^7^ CFU/ml at the last two time points (10 and 12hrs) **(Figure 2b)**. Surprisingly, biofilm formation ability remained unchanged until later stationary phase (10hrs) when biofilm biomass dropped by 25% compared to earlier time points **(Figure 2c-d)**. In other words, a substantial amount of biofilm biomass was still formed from planktonic culture in early stationary phase (8 hrs) when cellular proliferation is presumably diminishing, and the ability to form biofilm was lost only at 12hrs post infection. We also tested strains MAGAS5005 (serotype M1) and JRS4 (serotype M6) and, despite visible differences in biofilm morphology, observed that both exhibited strong biofilm phenotype when transferred at later stages of growth. By contrast, strain HSC5 (serotype M14) did not form robust biofilm after late exponential phase transfer **(Figure2 e-f)**. We next investigated the micro-phenotype of biofilms initiated from cultures at each growth phase. Consistent with CV assay results **(Figure 2c-d)**, SEM images showed slightly decreased surface coverage for biofilms initiated from 10 and 12hrs time points. In addition, the overall structure of biofilm initiated from 12 hr stationary phase cultures appeared less homogeneous tended to aggregate **(Figure 3a).** However, there were no major differences in biofilm structure or GAS chain organization between these biofilms **(Figure 3b).** As determined by OD600, CFU and biomass quantification, and imaging, these findings suggest that cell proliferation itself may not be crucial for biofilm formation in this static assay. Finally, we tested whether classic and transferred biofilms differ in antibiotic tolerance, a commonly used hallmark of biofilm [1, 5, 23]. We first determined the minimum inhibitory concentration (MIC), minimum bactericidal concentration (MBC) for planktonic cultures, and then compared the minimum biofilm eradication concentration (MBEC) for GAS JS95 biofilm upon exposure to penicillin. We observed that the MBEC of both types of biofilm were similar (differing by one double-dilution of the antibiotic) and far exceeded planktonic MIC and MBC values **(Figure S2).**

**Figure 2.**
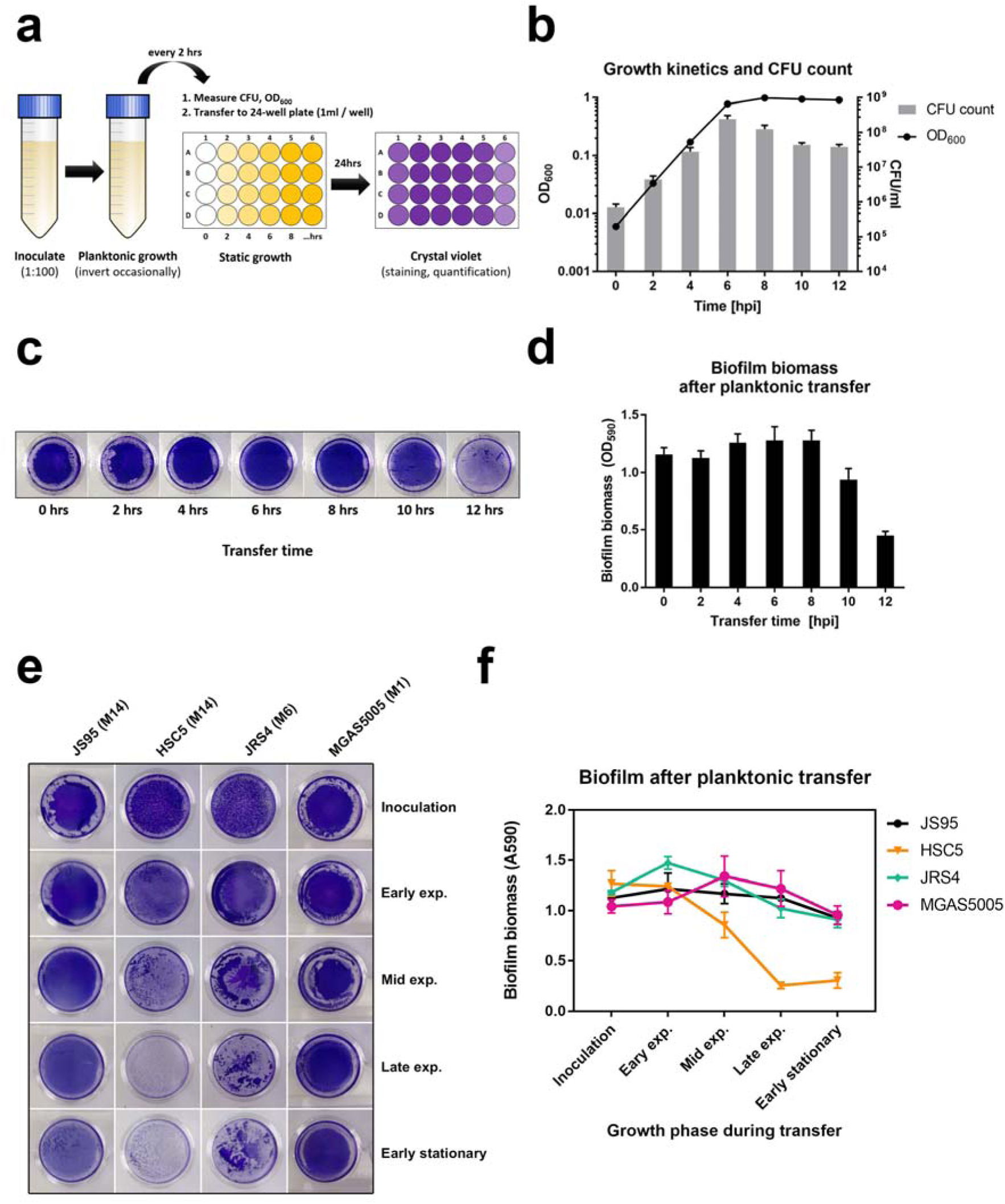
Planktonic transfer assay. **(a)** Assay scheme. Cells grown in planktonic culture in a 50ml conical tube were measured for CFU and OD600, then transferred to a separate 24-well plate. After 24hrs incubation, biofilm biomass was measured by crystal violet (CV) staining. **(b)** GD600 and CFU count at the indicated time-points during planktonic growth. **(c)** Images of biofilm stained with CV. **(d)** Biofilm biomass quantification by OD_590_ measurement of solubilized crystal violet. **(e)** Images and **(e)** quantification of biofilm formed by JS95, HSC5, JRS4 and MGAS5005 strains after planktonic transfer at various growth phases. Graphs show mean values +/- standard deviation.

**Figure 3.**
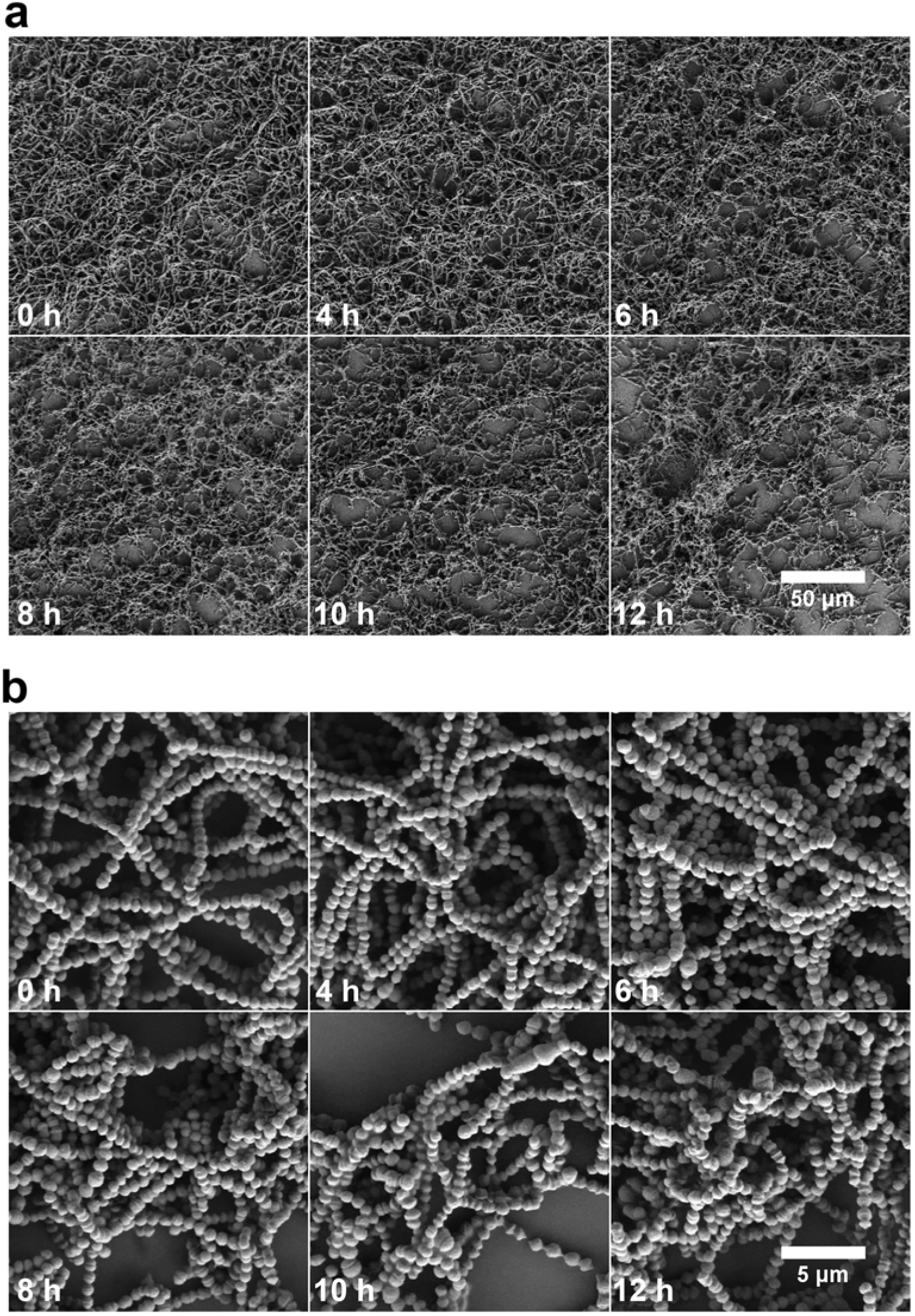
SEM images of GAS JS95 biofilm after planktonic transfer. Scanning electron micrographs (SEM) of biofilms from transferred planktonic cultures at the indicated timepoints. Biofilms were grown in 6-well polystyrene plates. Magnification **(a)** 500x and **(b)** 5000x.

### Cell proliferation is not necessary for adherent biomass accumulation

To determine whether cellular proliferation was indeed necessary for biofilm biomass accumulation, we treated GAS with the bacteriostatic antibiotic bacitracin. We first confirmed that bacitracin inhibits GAS proliferation at 4ug/ml, as reported previously [29]. Independent of the time of bacitracin addition (3, 4, 4.5, 5, 5.5, 6, 6.6, 7, 7.5 and 8 hours post inoculation), treatment with 4ug/ml bacitracin resulted in an OD600 plateau within approximately 1 hour, with no more than a 30% increase in turbidity after the time of treatment **(Figure 4a).** Therefore, we reasoned that after 2 hours of bacitracin treatment, there would be no further proliferation. Hence, we repeated the planktonic transfer assay, growing the planktonic culture for 8hrs (early stationary phase) and adding bacitracin at 4ug/ml for an additional 2hrs of incubation, and then transferring the non-proliferating planktonic cells to 24-well plate for static biofilm assay **(Figure 4b).** Crystal violet staining showed equally strong biofilm biomass regardless of bacitracin treatment **(Figure 4c-d).** Together these observations suggest that proliferation is not essential for biofilm biomass accumulation, which can arise from cells which are no longer dividing.

**Figure 4.**
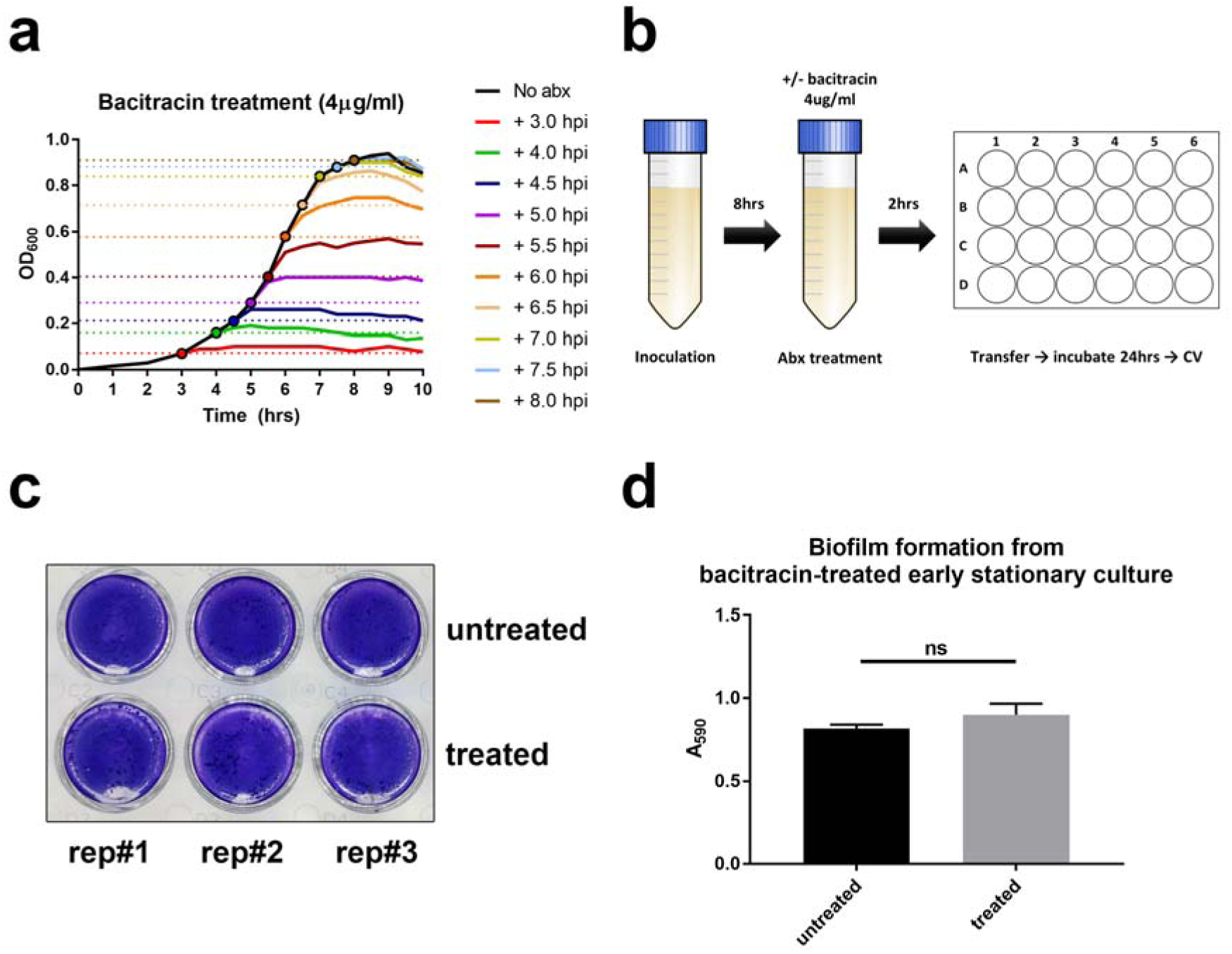
Bacitracin treated planktonic transfer assay. **(a)** Planktonic cultures were exposed to bacitracin (4µg/ml) at the time point indicated by the color-matched circle and growth inhibition was quantified by OD600 measurement. **(b)** Bacitracin treatment for planktonic transfer assay: planktonic culture was grown for 8hrs (to early stationary phase) prior to bacitracin treatment. After 2hrs incubation with antibiotic, cells from the non-dividing culture were transferred for static biofilm growth. **(c-d)** Biofilms were seeded from bacitracin treated and non-treated early stationary cells, and biomass was quantified by CV staining crystal violet after 24 hrs of growth.

### Planktonic transfer assay uncovers capsule-dependent phenotype differences, masked in classic CV assay

Cho and Caparon previously showed that a GAS mutant deficient in capsule production was able to form biofilm in static conditions as well as the isogenic parental wild-type strain, but was unable to develop biofilm in flow biofilm conditions [19]. We therefore hypothesized that some of the phenotypic inconsistencies between these assays could be dissected using the planktonic transfer assay established in this study. We performed planktonic transfer assay at 1-hour intervals for a total of 10 hrs, comparing wild type JS95 and an isogenic *hasA* mutant deficient for capsule production [17]. **(Figure 5a).** WT GAS JS95 biofilms inoculated from all growth phases, including early stationary phase, resulted in dense biomass accumulation, as was observed above **(Figure 2c, d).** In addition, we observed visually distinct CV staining for early transfer times, whereby the *hasA* mutant biofilm appeared more robust, although CV quantification did not reveal significant differences **(Figure 5a-b).** By contrast, the *hasA* mutant was unable to accumulate biomass when biofilms were initiated from mid exponential cultures onward **(Figure 5a-b).** Interestingly, both strains (WT and *hasA* mutant) were unable to form biofilm in Calgary biofilm device (CBD), in which sedimentation of cells that might contribute to biofilm formation is excluded due to the inverted device geometry **(Figure S3)** [7, 30, 31]. Strong biofilm was, however, observed in *P. aeruginosa* which served as a positive control [9]. To explore the contribution of adhesins to biofilm formation by each mechanism, we used simple adhesion assay in which planktonic cultures at different growth stages were normalized to the same optical density, incubated in a 24 well plate for 30min in 37C, non-adherent cells removed by washing, and the adherent cells stained using crystal violet. We observed that adhesion of the capsule mutant was similar at all tested growth phases **(Figure 5c).** Adherence of the wild type strain increased gradually from low levels in early exponential growth, to late stationary phase where it reached the same level as the capsule mutant. We confirmed that, as in other GAS strains, *hasA* expression peaks at mid-exponential growth, then decreases rapidly **(Figure 5d)** [32, 33]. Together with the fact that capsule shedding is associated with cessation of its synthesis [34], these data provide further evidence that the capsule hinders adhesin-mediated surface attachment. In the same adhesion assay we tested JS95 mutant unable to produce M-protein, which is a well-studied GAS virulence factor, also shown to be crucial for biofilm formation [19]. By contrast, a mutant in M-protein, was non-adhesive until the late stationary phase (overnight culture), at when adherence reached approximately half the level of WT at the same growth phase, showing that surface adhesion is not solely dependent on M-protein. Finally, we tested whether exogenous hyaluronidase treatment of WT biofilms could phenocopy *hasA*^-^ biofilms. Indeed, capsule removal by hyaluronidase slightly enhanced classic WT biofilm formation and significantly reduced biofilm formed from cultures transferred at 8hpi **(Figure 5e).** At the same time, hyaluronidase presence had little to no effect on *hasA*^-^ mutant. These data suggest that the capsule contributes differently to biofilms depending on growth stage: it inhibits biofilm formation during early exponential growth by masking surface adhesins, but promotes biofilm formation when it is initiated from later phase cultures.

**Figure 5.**
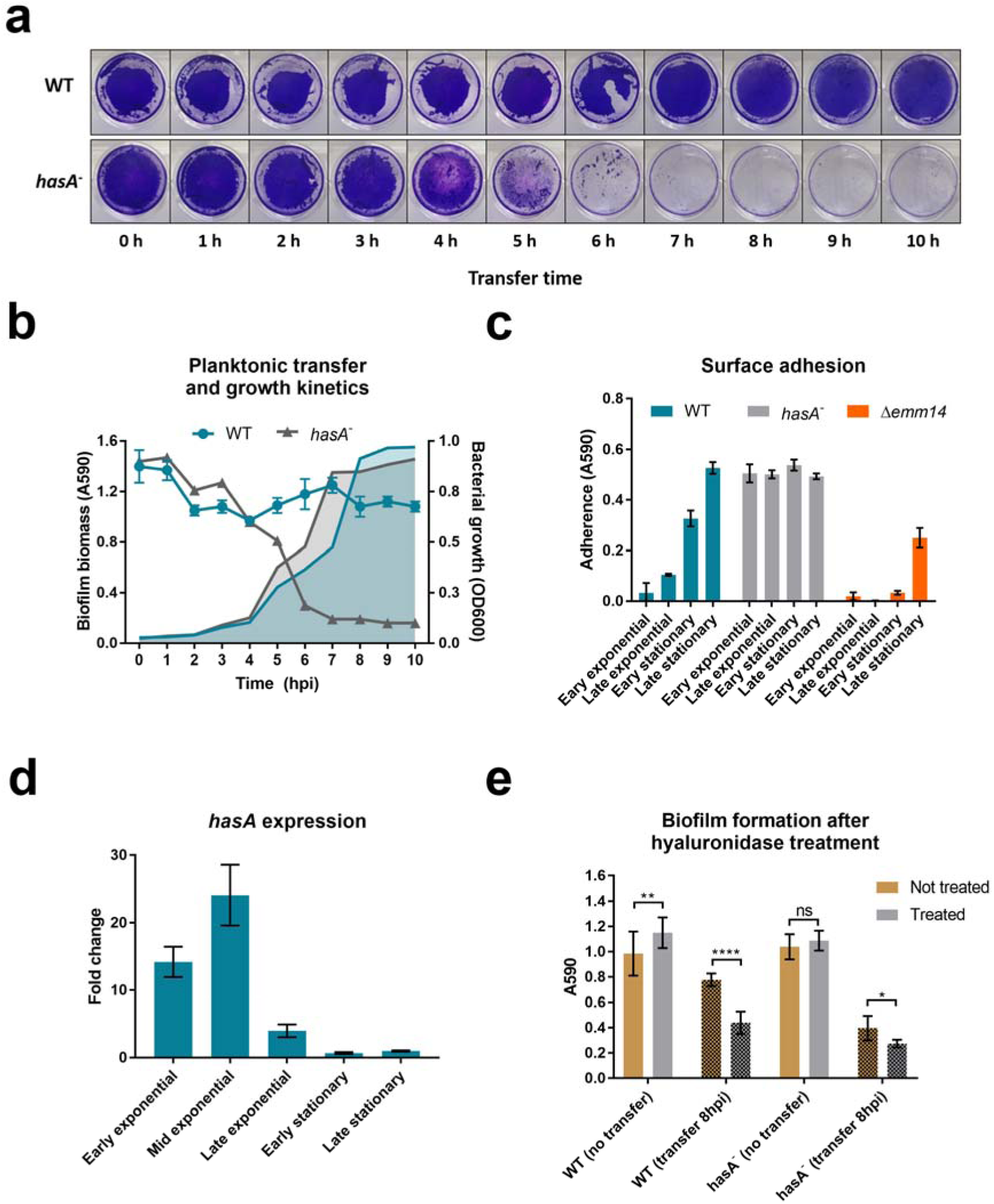
Planktonic transfer assay with capsule deficient GAS mutant. **(a)** CV stained WT and *hasA*^-^ biofilms transferred at the indicated time points. **(b)** Planktonic bacterial growth (OD600) and biofilm biomass quantified by CV staining (A590) at the time of transfer. **(c)** Adherence of JS95 planktonic cultures (normalized to the same OD600) to polystyrene, derived from the indicated growth phases. **(d)** *hasA*^-^ expression at different phases of growth in THY + 0.5% glucose, relative to late stationary (15hpi) culture. **(e)** Hyaluronidase treatment (50µg/ml) of classic biofilms transferred at 8hrs (early stationary phase). Graphs show mean values +/- standard deviation.

## DISCUSSION

In this study we investigated biofilm formation by the clinical GAS strain JS95 (M14 serotype), isolated from an NF patient [18]. We confirmed that it forms biofilm in static conditions, showing dense three-dimensional biofilm structures of chaining cocci characteristic for many of the GAS strains. Although GAS has been shown to produce a biofilm matrix of extracellular polymeric substances (EPS), composed primarily of L-glucose and D-mannose [35], we were unable to observe an extracellular EPS matrix using microscopy for either WT or *hasA*^-^ stains.

Strain-associated differences in GAS biofilm formation have been reported. To address whether strain-independent factors might contribute to biofilm formation, we were motivated to step back and more carefully assess how GAS strain JS95 biofilm is formed using the common static biofilm assay. Using the planktonic transfer assay described here, we observed that GAS cells from planktonic cultures ranging from early exponential phase to early stationary phase can seed robust biofilm formation and biomass accumulation. We only observed reduced biomass accumulation when we seeded biofilms with cells from 12 hr late stationary phase cultures. This phenotype was not strain specific, because GAS strains of different serotypes (JRS4, MGAS5005) displayed a similar ability to form biofilm from late exponential / early stationary planktonic cultures. Moreover, bacitracin inhibition of cell proliferation did not prevent biomass accumulation in these conditions. Together these data suggested the existence of two parallel mechanisms of static biofilm formation. 1) In the first, “classic” mechanism, a variety of adhesins including M and M-like proteins facilitate initial surface attachment, followed by microcolony formation, cell proliferation, biofilm matrix production, and biofilm maturation [9, 36]. 2) In an alternative mechanism, planktonically growing cells eventually sediment and attach to the surface, in a process that is enhanced by GAS capsule, leading to biofilm formation **(Figure 6a).** In support of this model, biofilm development from cells seeded from early exponential phase culture (2h post inoculation) showed typical steps of cell attachment to surface (2-3h post transfer), microcolony formation (4-5h post transfer), and dense biofilm formed as early as 6 h post transfer (8h post inoculation). By contrast, cells seeded from late exponential phase culture (6h post inoculation) showed some level of initial surface attachment 2 hours post transfer, and a rapid increase of biofilm density at later time points **(Figure 6b).** Both situations led to stable and dense GAS JS95 biofilm formation. Importantly, the second, alternate mechanism for biofilm formation could only be uncovered using the planktonic transfer assay, where sedimentation of later growth phase cultures can promote biofilm biomass accumulation. By contrast, biofilm biomass accumulation in the classic static assay is likely a result of both mechanisms (early attachment and proliferation of microcolonies, as well as sedimentation of later growth phase bacteria). A similar suggestion that sedimented cells could contribute to biofilm biomass led to the development of the Calgary biofilm device (CBD) eliminate the possibility of sedimentation [7, 30, 31]. However, GAS-JS95 did not form biofilms on CBD, further suggesting a strong contribution of the alternate mechanism of GAS biofilm formation.

**Figure 6.**
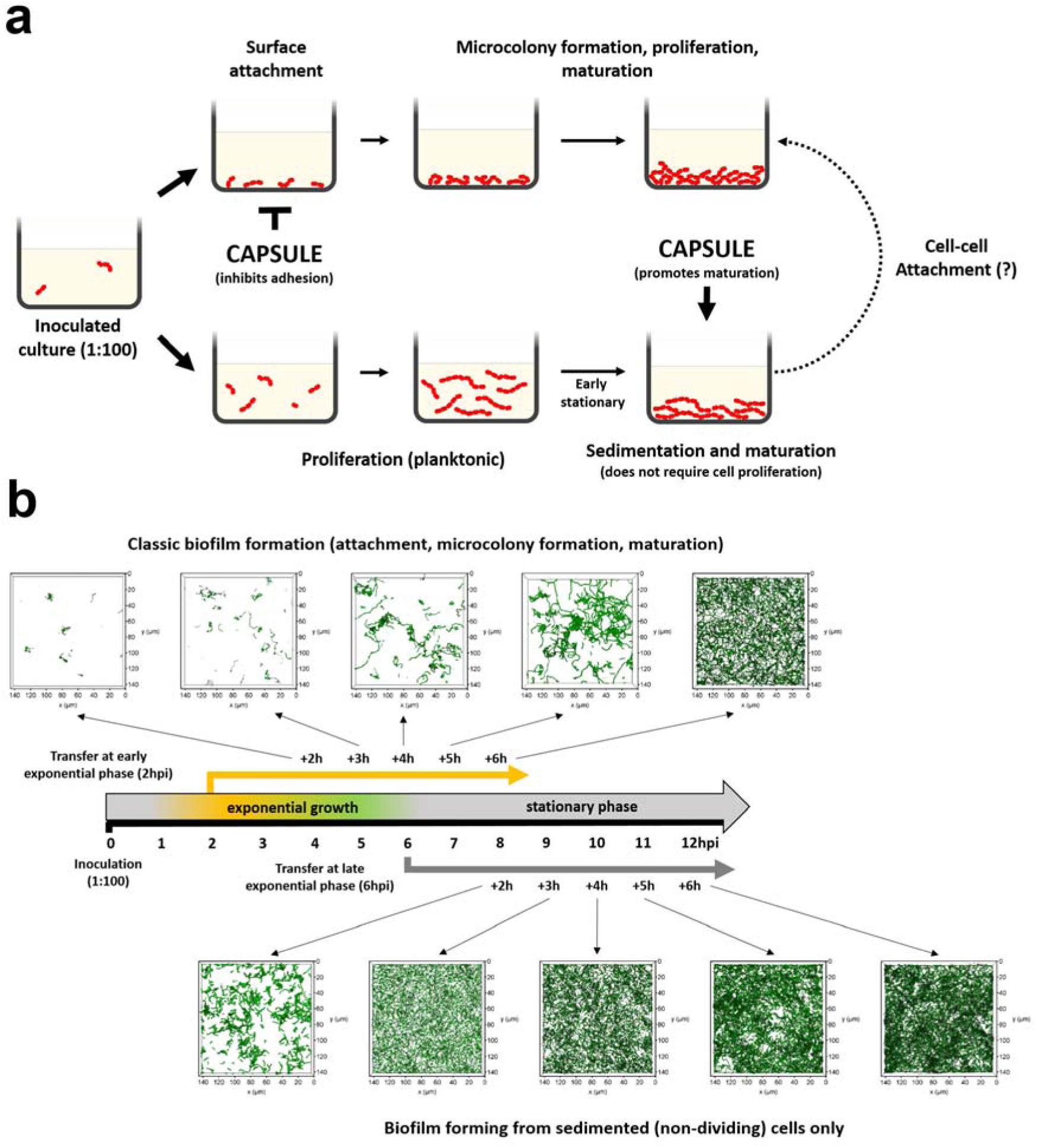
Model for static GAS biofilm formation. **(a)** In the classic mechanism for biofilm formation, surface attachment is followed by microcolony formation, cell proliferation, EPS production, and biofilm maturation. In the alternate pathway for biofilm formation, planktonic cells sediment when they reach a critical cell density, attach to a surface, and become fixed into biofilm structure. Both mechanisms are likely occur in the widely used static biofilm assay. **(b)** Time course showing biofilm development from culture transferred at early and late exponential phases. Images of biofilm developing from early exponential culture show typical steps of initial attachment (2-3h of static incubation), microcolony formation (4-5h) and stable biofilm (6h). Despite higher cell density, transfer of late exponential phase cells result in moderate initial attachment (2h of static incubation) followed by rapid increase biofilm formation a later time points, supporting the proposed alternate sedimentation mechanism of biofilm initiation. CLSM images of Syto9 stained biofilms are rendered as volume projection.

Although many studies have demonstrated a strong link between capsule production and GAS virulence, the role for capsule in GAS biofilm formation has been unclear [36-38]. Capsule production is highly regulated during GAS growth, with minimal expression during stationary phase (enabling initial adherence in a classic biofilm model), and peak expression during exponential phase (supporting an alternate, capsule-dependent model for biofilm formation) [32-34], and the same pattern can be observed in JS95 strain. Cleary et al. noted that encapsulated cells grew in a highly aggregated state that can be disrupted by hyaluronidase treatment [39]. These capsule-associated aggregates were originally described as a protective mechanism against oxidative stress, equally consistent with the protective and environmental stress-tolerant state associated with biofilms. šmitran et al. demonstrated that enzymatic removal of capsule prior to biofilm initiation improved static biofilm formation by most GAS isolates [40], suggesting that the capsule masks biofilm-associated surface adhesins, as has been demonstrated M-protein mediated GAS attachment to keratinocytes [41]. In the context of our revised model for GAS biofilm formation, initial capsule removal would be expected to promote classic biofilm formation, where initial adhesion is essential for biofilm development. Consistent with this, we observed altered early stage biofilm formation whereby the *hasA* mutant biofilm appeared more robust, although this was not quantifiable by CV staining. The importance of capsule in later stages of biofilm formation has been shown by Cho and Caparon, who demonstrated that a capsule mutant was unaffected in surface adhesion, but was unable to form biofilm in a flow cell [19]. However, both WT and the capsule mutant were fully capable of forming biofilm in the classic static biofilm assay [19]. The ability of the planktonic transfer assay to detect different mechanisms of biofilm initiation enabled us to further dissect the contribution of capsule to biofilm formation. Our finding that capsule mutants are attenuated only for sedimentation-mediated, microcolony-independent static biofilms may suggest that GAS sedimentation characteristics may be important in the maturation of flow cell GAS biofilms. Collectively, these findings suggest that capsule may limit initial surface adherence and promotes sedimentation-mediated biofilm maturation. The ability of the planktonic transfer assay to dissect he contribution of growth stage specific factors may be widely applicable to many other GAS biofilm factors as well.

In summary, we showed that different mechanisms contribute to static GAS biofilm formation: 1) classic surface adhesion followed by microcolony formation and biofilm maturation; and 2) sedimentation and attachment of dense planktonic bacteria to a surface followed by biofilm maturation. We showed that capsule may differentially contribute to each of these mechanisms. Separation of these two mechanisms might help to uncover phenotypes otherwise masked in static biofilm assays, enabling a better understanding of the mechanisms of GAS biofilm formation, and ultimately may inform how biofilm relates to GAS virulence.

## MATERIALS AND METHODS

### Bacterial culture, planktonic transfer assay, and biofilm assay

GAS strain JS95, an M14 serotype isolate from a NF patient [18], or a *hasA* mutant in the JS95 background, as well as strain JRS4 (M6 serotype), HSC5 (M14 serotype) and MGAS5005 (M1 serotype), were grown at 37°C overnight (16hrs) in Todd-Hewitt liquid medium (Sigma Aldrich) supplemented with 0.2% yeast extract (Becton, Dickinson) (THY) prior to all assays. Overnight cultures were inoculated 1:100 into fresh THY medium supplemented with 0.5% glucose in a 50ml tube, and incubated at 37°C, with 5% CO_2_ and a loose tube cap. Cultures were mixed by inverting the tube three times at 30 min intervals to prevent sedimentation. At the desired time points, 1 or 3 ml of planktonic culture was transferred into a well of a 24 well (for crystal violet staining) or 6 well (for SEM) polystyrene plate (Corning, New York, USA), respectively. For confocal laser scanning microscopy (CLSM), 3ml of culture was transferred to a 35mm Ibidi imaging dish (Ibidi, Munich, Germany), which allow high resolution CLSM imaging. Although some differences between substrates might influence biofilm formation, we did not observe a significant impact on biofilm formation between polystyrene and the Ibidi surface. Plates or Ibidi dishes were then incubated without agitation for 24 hrs at 37°C with 5% CO_2_. At the time of transfer optical density at 600nm was measured in a 1cm cuvette by UVmini-1240 UV-Vis spectrophotometer (Shimadzu, Japan) and bacterial viability was quantified by serial colony forming unit (CFU) enumeration. Prior to CFU plating, cultures were centrifuged for 2 min at 15,000 rcf using a table top Eppendorf 5424 centrifuge (Eppendorf, Germany) to disrupt chains (confirmed by bright field microscopy). To inhibit bacterial proliferation, bacitracin (Sigma Aldrich) was added at 4 µg/ml for the times indicated.

### Biofilm quantification

After 24 hrs of growth, non-adherent cells were washed gently with PBS. Biofilms were stained with 0.1% crystal violet (CV) for 15min, and the excess CV was removed by a subsequent PBS wash. Stained biofilms were first imaged using a Huawei Mate 20 Pro camera (Huawei, China) and dissolved in 96% ethanol. Prior to OD measurement, samples were diluted to ensure a linear reading range. Absorbance was measured at 590nm using Tecan M200 spectrophotometer (Tecan AG, Männedorf, Switzerland).

### MIC, MBC and MBEC assay

Standard protocols with slight modifications were used to determine the MIC, MBC and MBEC values [42, 43]. Briefly, to determine the MIC, subcultures (inoculated 1:100) were grown overnight in a 96well plate in THY with serially diluted antibiotic. Turbidity was assessed using a plate reader (Tecan AG, Männedorf, Switzerland) at 600nm to determine the minimal antibiotic concentration that inhibited bacterial growth. Cultures were then subcultured onto THY agar plates to determine the MBC. To determine the MBEC, biofilms were prepared by inoculating THY + 0.5% glucose 1:100 from an overnight culture and transferring (1ml/well) to 24-well plate (referred in text as “classic” biofilm) or incubating inoculated medium in 50ml conical tube for 8 hours (until early stationary phase) and inverting occasionally (to maintain cells in a planktonic state), followed by transfer to 24-well plate (referred in text as “transferred” biofilm”). Both types of biofilm were incubated 16 hours post inoculation, then washed gently 2 times with PBS, exposed to THY containing various concentration of penicillin, incubated 1hr, washed again 2 times with PBS, and incubated overnight in fresh THY. Turbidity was then measured using a plate reader at 600nm and the lowest concentration of antibiotic giving no turbidity was defined as MBEC. All incubations were at 37°C.

### Hyaluronidase treatment

“Classic” and “transferred” biofilms were prepared as described for the MBEC assay, except both types of biofilms were incubated for 24 hours in 37°C and inoculated 1:100 medium was supplemented with 50µg/ml hyaluronidase (cat#H3506, Sigma-Aldrich). Following incubation, biofilm was quantified as described above.

### Surface adhesion

Fresh THY + 0.5% glucose was inoculated 1:100 with an overnight culture grown in THY and incubated at 37°C in a conical tube until the desired growth phase was reached: early exponential (OD600 = 0.2), late exponential (OD600 = 0.8), early stationary (OD600 reaching plateau, around 8h), late stationary (16h). Cultures were then spun for 10min at 8000rcf, resuspended and normalized to OD600 = 1.0 in PBS, transferred to 24-well plate (1ml/well), incubated for 30min at 37 ° C, washed twice with PBS, then stained with 0.1% crystal violet for 10min and washed again with PBS. Crystal violet was then solubilized in 96% ethanol and quantified at 590nm using a Tecan M200 spectrophotometer (Tecan AG, Männedorf, Switzerland).

### RNA extraction and RT-qPCR

Bacteria grown in THY + 0.5% glucose were harvested by centrifugation for 1min at 10,000rcf at various growth stages: early exponential (OD600 = 0.2), mid exponential (OD600 = 0.5), late exponential (OD600 = 0.8), early stationary (OD600 reaching plateau, around 8h), late stationary (16h). RNA was isolated using Direct-zol RNA MiniPrep (Zymo Research, USA), and DNAse treated with Turbo DNA-free kit (Ambion, USA). RNA concentration and DNA absece was assessed by Qubit 2.0 (Invitrogen, USA) and the integrity was determined by Tape Station (Agilent Technologies, USA). Samples with RIN > 7 and DNA contamination <10% were used for cDNA synthesis using SuperScript III first-strand synthesis kit (Invitrogen). RT-qPCR was performed using 2xSYBR FAST qPCR universal MasterMix kit (Kappa Biosystems, USA). Gyrase A (*gyrA*) was used as an endogenous control [44]. The following primers were used (5’ > 3’): *hasA:* (FWD) AGGACGCACTGTCTACCAATC; (REV) GTCCATAAGGCAACGATGGGA; *gyrA:* (FWD) CAACGCACGTAAGGAAGAAA; (REV) CGCTTGTCAAAACGACGTTA.

### Confocal laser scanning microscopy

Biofilm in 35mm a Ibidi dish was washed gently in PBS (twice), then fixed with 4% paraformaldehyde for 10min, washed again with PBS, and stained with the indicated dyes according to manufactrer protocol and washed again with PBS. Matrix staining was performed using WGA-AF633 at 50 μg/ml. Hoechst33342 and Syto9 were used to stain DNA of all cells. Propidium iodide was used to stain DNA of membrane-compromised cells, as it cannot cross the membrane of live cells. For live-dead staining, cells were not fixed with paraformaldehyde. Microscopy was performed using a Zeiss LSM780 confocal microscope, equipped with 20× NA0.80, 63× NA1.40 (oil) or 100× NA1.46 (oil) lens. Collected Z-stacks were projected as volume or maximum projection using FIJI distribution of ImageJ (NIH, Maryland, USA).

### Scanning Electron Microscopy

Biofilms in 6 well plates were washed 3x with 0.1M PB, fixed overnight in 4°C with 2.5% glutaraldehyde (Agar Scientific), post fixed with 1% OsO4, dehydrated in an ethanol gradient, and finally dried using HMDS (Sigma Aldrich). After platinum coating, samples were imaged using Jeol 7610F instrument (Jeol, USA).

## ACKNOWLEDGMENTS

We would like to thank Dr. Emanuel Hanski for providing bacterial strains used in this study, Alicia Tan and Ho Foo Kiong for help with experiments, and the members of the Kline lab for critical reading of this manuscript. The authors acknowledge financial support from the Singapore Centre for Environmental Life Sciences Engineering (SCELSE), whose research is supported by the National Research Foundation Singapore, Ministry of Education, Nanyang Technological University and National University of Singapore, under its Research Centre of Excellence Programme. This work was also supported by the National Medical Research Council under its Clinical Basic Research Grant (NMRC/CBRG/0086/2015).

## SUPPLEMENTARY FIGURES

**Figure S1.**
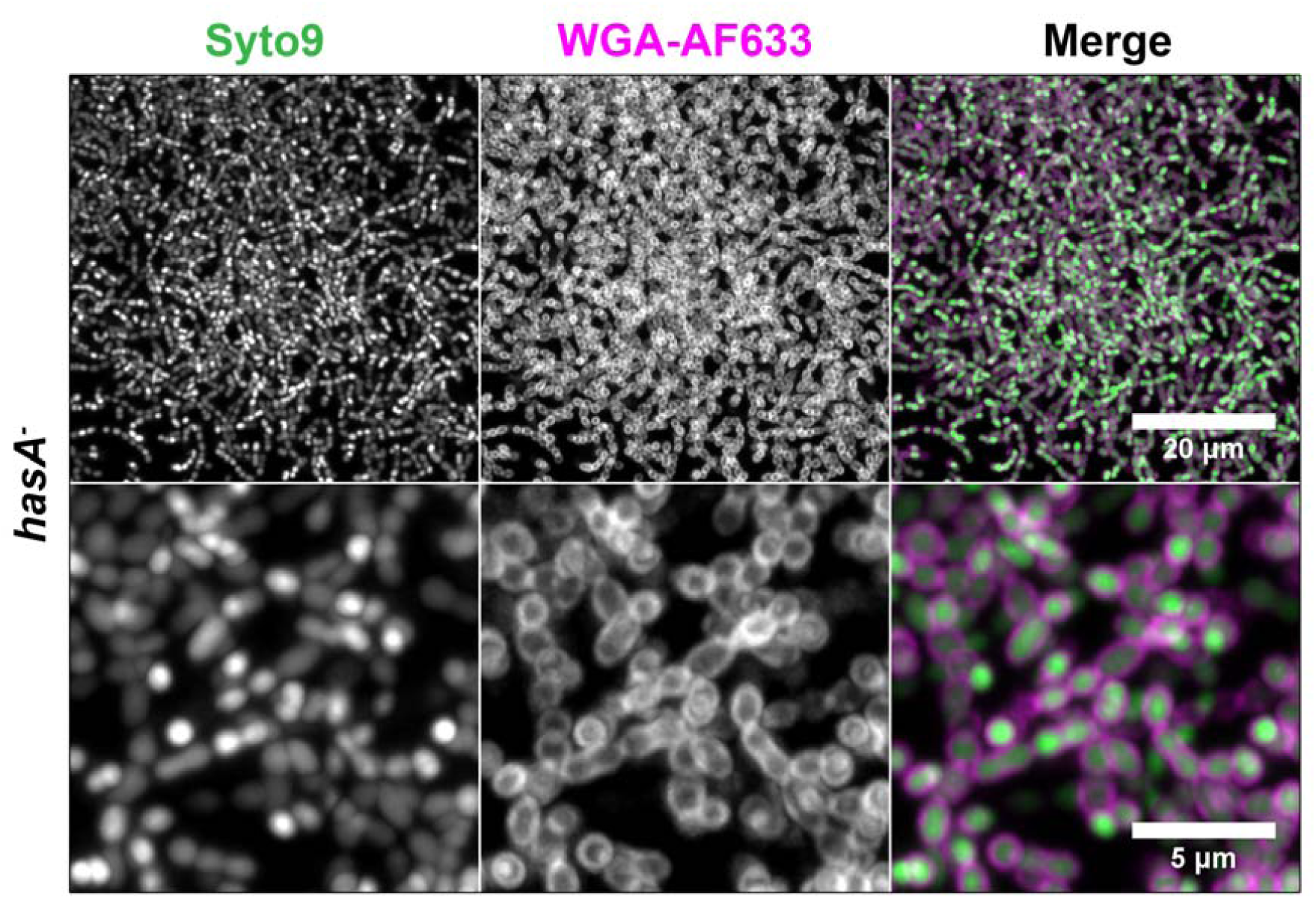
WGA staining. Maximum projection of JS95 *hasA-* biofilm Z-stacks stained with Syto9 (dsDNA) and WGA-Alexa Fluor 633 (carbohydrate/EPS). Positive staining of capsule null mutant confirms that WGA-AF633 signal does not originate from non-specific capsule staining. As in WT, the matrix components visible only at the cell surface but not in extracellular space.

**Figure S2.**
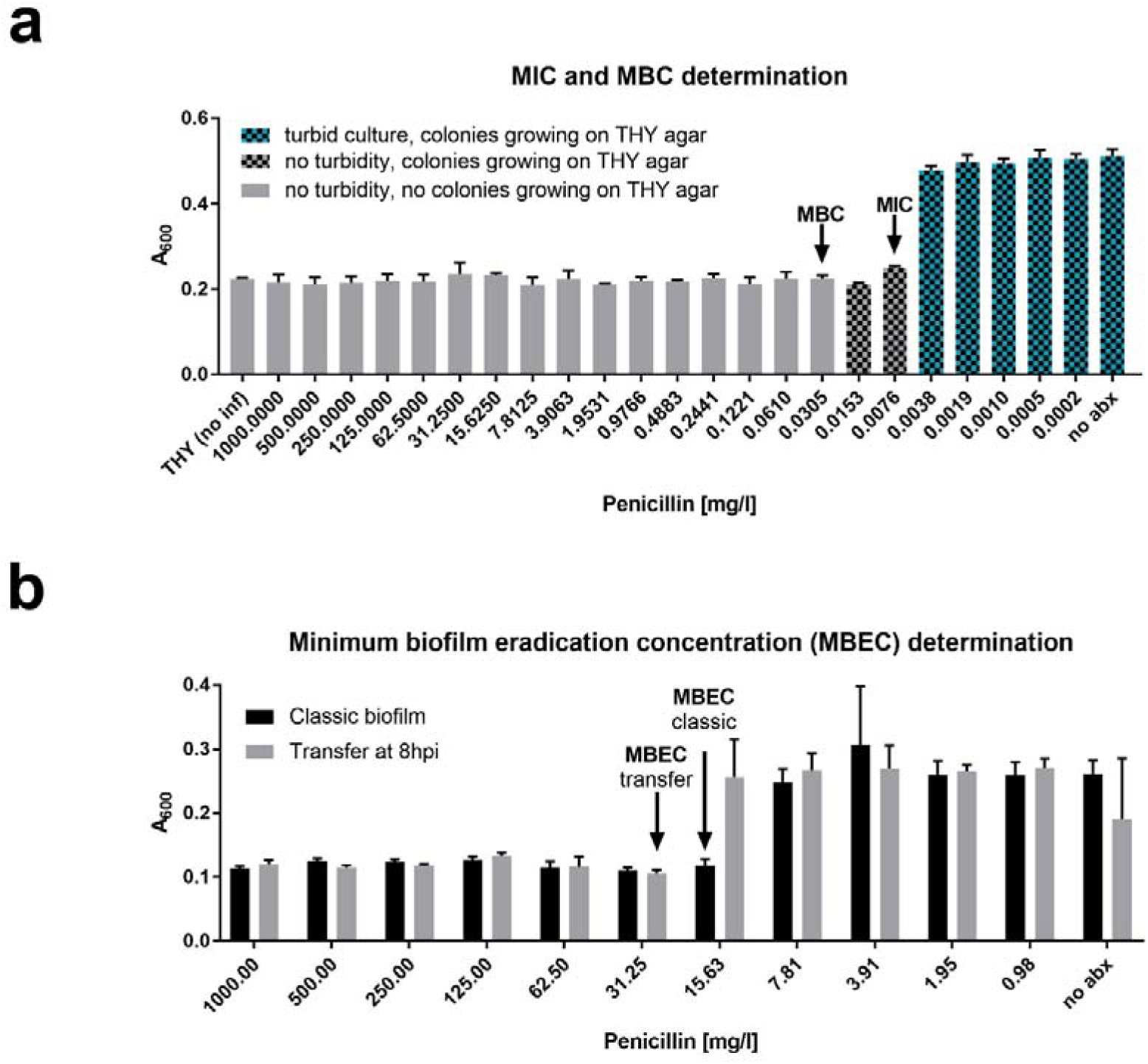
**(a)** Minimal inhibitory concentration (MIC) and minimal bactericidal concentration (MBC) determination. Bars indicate absorbance at 600nm of cultures incubated overnight with the indicated concentrations of penicillin. Grid patterned bars indicate bacterial growth when plated on THY agar. **(b)** Minimal biofilm eradication concentration (MBEC) estimated by challenging classic and transferred biofilms with penicillin for 1hr, followed by PBS wash and overnight incubation with rescue medium. Bars indicate absorbance at 600nm.

**Figure S3.**
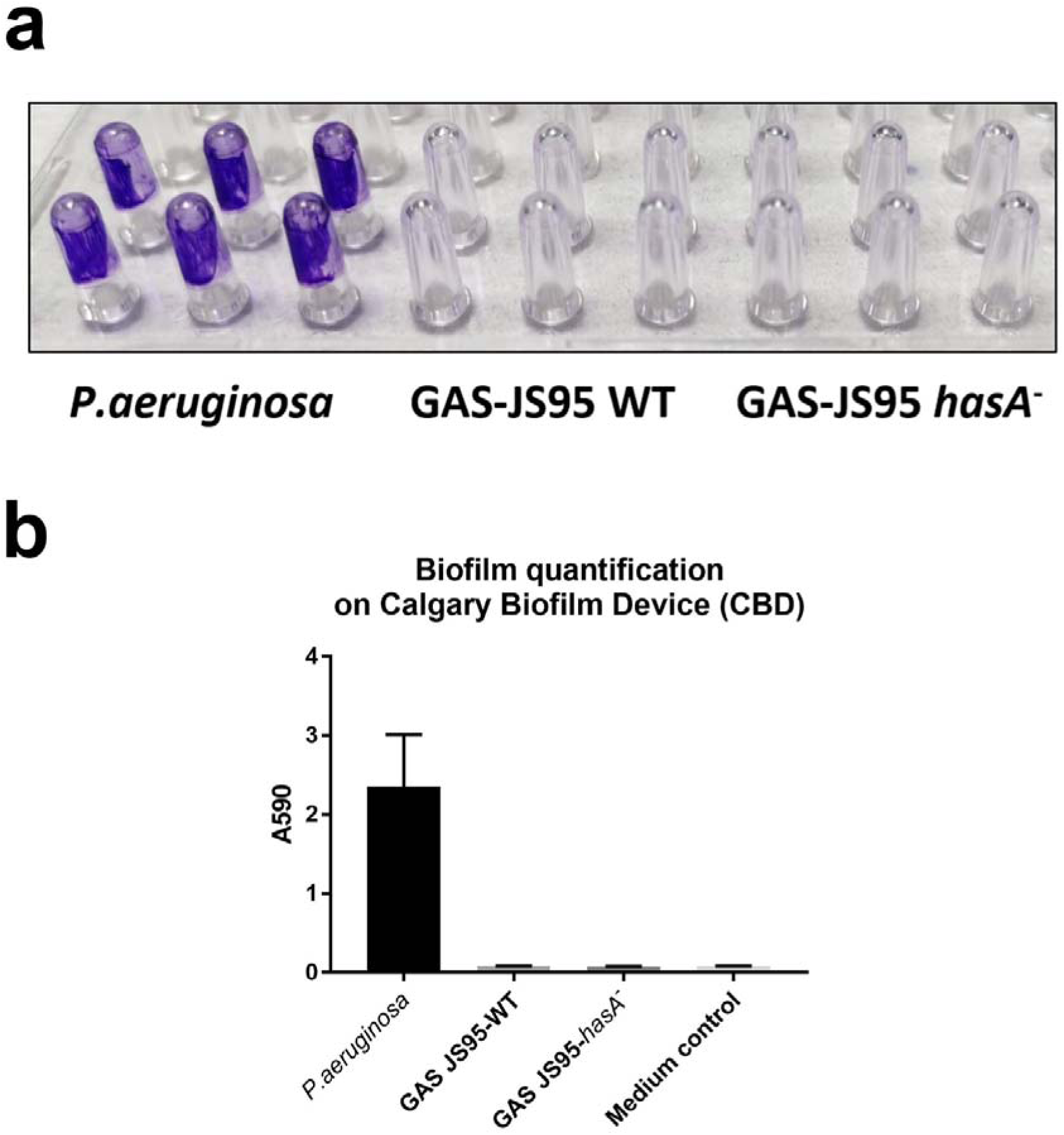
**(a)** Biofilm grown on pegs of Calgary biofilm device (CBD), washed and stained with crystal violet. *P. aeruginosa,* a classic biofilm former, was used as positive biofilm control. There is no visible growth of JS95 wild type (WT) as well as capsule mutant (*hasA*-) **(b)** Quantification of CBD biofilm after solubilisation in ethanol.

